# Some Mendelian Disorders could be fixed with a pill? A Structural Bioinformatic investigation

**DOI:** 10.1101/2022.10.20.513070

**Authors:** Pietro Bongini, Daniele Iovinelli, Alfonso Trezza, Monica Bianchini, Ottavia Spiga, Simone Gardini, Neri Niccolai

## Abstract

It has been recently suggested that amino acid replacements with Gly can modify the shape of protein surfaces and, hence, protein dynamics and functions. We have browsed ClinVar, the database of all the reported variants of clinical relevance, to identify all the proteins having missense X/Gly mutations that determine Mendelian disorders. We have found 959 benign and 875 pathogenic X/Gly substitutions. Pathogenicity origins were initially searched in the distribution profiles of replaced amino acids. These profiles indicate that Mendelian disorders including Gly-replacements arise mainly from substitutions of amino acids bearing bulky hydrophobic side chains, thus reducing protein core stability. In the case mutated proteins were structurally defined, we could give a deeper insight into pathogenicity mechanisms, checking whether Gly-mutations altered protein shapes, modifying water surface dynamics and, hence, the physiological protein-protein interaction processes. In several cases, indeed, we have found that pathological Gly-mutants present additional surface pockets, suggesting that the new pockets could be the target of a pharmacological strategy for Mendelian disorder remediation.

**Highlights:**

- ClinVar has been scanned to find signals for pathogenicity due to X/Gly mutations
- Pathogenicity origins of X/Gly replacements have been structurally analyzed
- X/Gly mutations can create protein surface pockets with binding capabilities
- Gly-formed new protein binding sites can be the target for Mendelian disorder cures
- AI procedures will expand the search for structural damages due to X/Gly mutations

**Graphical abstract:** 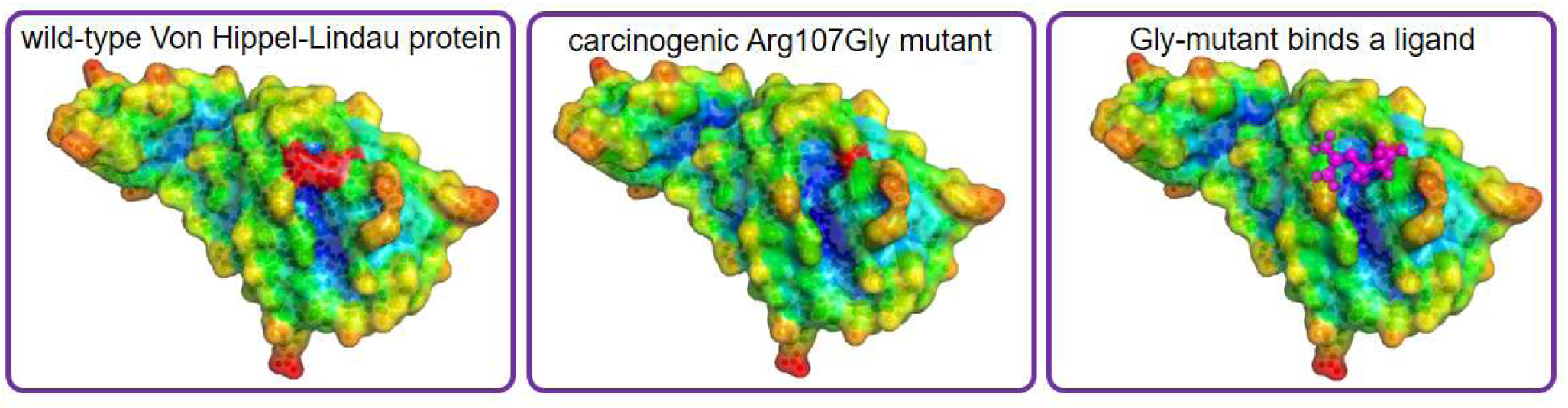

## 1. Introduction

In the genomic era, personalized medicine is taking advantage of the continuous progress in the development of fast and accurate procedures for genome annotation and variant classification [1]. Thus, we can expect that an increasing number of single-gene disorders that are due to variants in a specific gene, also referred to as Mendelian disorders, will be characterized and listed in genome variant databanks, such as OMIM [2], HuVarBase [3], or ClinVar [4].

Several therapeutic strategies for the treatment of Mendelian disorders are currently in use (see [5] for a review), including replacement of deficient gene products, manipulation of gene expression and pre-mRNA splicing, and compensation for functional deficits through new applications of FDA-approved drugs. The latter approach is particularly promising in the artificial intelligence-driven search for drugs that could, at least in part, remediate Mendelian disorders [6].

Among all Mendelian disorders, single nucleotide variants causing pathogenic missense mutations can be analyzed in detail, particularly in the case that structural information is available for the mutated protein. Structural analysis of mutants, indeed, can indicate the type of damage induced by the mutation, suggesting how remediation can be pharmacologically achieved to restore original functions.

The large array of resolved protein structures that are present in the Protein Data Bank, PDB [7], can be used to correlate pathogenicity with structural changes in mutants [8]. Thus, by scanning PDB content to find the structural features of proteins involved in missense mutations and included in genome variant databanks, we can obtain powerful clues for developing new drugs or repurposing the already approved ones to cure Mendelian disorders.

Recently, it has been shown that replacements of amino acids with Gly may determine the formation of new surface pockets, modifying protein surface structure and dynamics and, hence, protein function [9]. Thus, we explored the possibility that the replacement of a generic amino acid X with Gly, would yield an X/Gly missense mutation which is responsible for a Mendelian disorder. In the present investigation, we describe how X/Gly missense mutations can modify the structure of a protein and, particularly, its surface, leading to the formation of new pockets. The Structural Bioinformatics procedures we have used here, are simple and they rapidly predict the existence of Gly-mutant sites that can be the target of small molecules, whose binding could restore the original surface dynamics and, hence, the physiological protein function.

## 2. Methods

ClinVar is a very large public database of reported associations between human variants and phenotypes [4]. To select the relevant information for the present study, we applied the following filters through the web interface of ClinVar: “type of variation” = “single nucleotide variant”; “molecular consequence” = “missense”; “review status” = “at least one star”. Then, by applying the “clinical significance” filter, we extracted two separate mutation data sets with “benign” and “pathogenic” filter values. Finally, we selected only those mutations that lead to X/Gly substitutions, in both benign and pathogenic data sets.

To explore the structural consequences of Gly-mutations, we had to take into account only proteins having PDB reference files. Then, we could use *Gly-pipe*, a software pipeline for the prediction and identification of potentially druggable surface pockets induced by glycine substitutions [10]. We checked whether the sequence coordinate of Gly-mutation given by each ClinVar entry corresponded in the PDB file either to the wild-type amino acid or Gly. All the PDB entries not fulfilling this requirement were discarded. The first step of the *Gly-pipe* routine consists in substituting the wild-type amino acid with Gly (direct substitution) or Gly with the wild-type amino acid (inverse substitution), through the PyMOL-based substitution module [11]. Either way of applying the substitution results in a pair of structures: one wild-type and one mutant structure.

Fpocket [12] analyzes the latter two structures to compare their protein surfaces for detecting the appearance of a new cavity near the mutated amino acid or the widening of a preexistent pocket. Both of these events are linked to the estimation of pocket druggability score (DS) through the neural-network-based module of *Gly-pipe*. We have used Open source PyMOL v. 1.7.1.0 for structural data analysis and to generate rational Gly mutants. For refining PyMOL mutation procedures, we used the *Optimize* PyMOL plugin [13]. Protein surfaces were colored according to atom depths as calculated with the SADIC algorithm [14], by using the freely downloadable software at http://www.sbl.unisi.it.

## 3. Results

We have searched for X/Gly missense mutations among all the items that were collected in the ClinVar variant databank. As of July 4, 2022, ClinVar contained 1,546,548 human genomic variants, 594,188 of which yield missense mutations. ClinVar reported 33,919 of the latter as benign and 27,521 of them as pathogenic variants. By parsing the information contained in the latter ClinVar items, we have found 959 benign and 875 pathogenic X/Gly substitutions. The fact that the number of these substitutions is similarly lower than the average value which should be expected for the 20 naturally occurring amino acids, *i*.*e*. respectively -25.0 % and -23.5%, reflects the limited damage that is caused by the replacement of Gly with its small and uncharged side chain. It must be noted also, that all the eight possible X/Gly substitutions, arising from a single-nucleotide codon change, are present in the latter 1.834 Gly-mutants, even though at very different extents.

From the amino acid profiles reported in Fig. 1, several features are apparent: Gly-substitutions of amino acids with small side chains, such as Ala and Ser, determine very limited pathogenicity, indicating that the volume reduction of Gly side chain is well tolerated in the mutant structure. This is not, of course, the case of Cys, whose determinant role in structural stability through the formation of cystine-bridges is well known. Furthermore, damages in protein folding are ascribed to Gly replacements to amino acids with bulky hydrophobic side chains. Thus, Val/Gly and Trp/Gly mutations most likely interfere with the proper formation of the protein folding nucleus and, hence, with the achievement of the bioactive protein structure [15]. In this respect, pathogenicity enhancement due to Trp/Gly substitutions appears to be even more critical than the Cys/Gly ones, as the benign/pathogenic replacement ratios are respectively 14% and 31%. In principle, the structural consequences of X/Gly mutations could be explored case by case for each of the mutated proteins reviewed by ClinVar, as algorithms have been proposed for predicting the effects of a mutation on the protein folding nucleus formation [16], but it will not be done in this report. Fig. 1, highlights the frequent occurrence of Gly replacements with Arg, Asp and Glu with pathogenic and benign similar impacts. The latter amino acids, all with charged side chains, are usually found in the outer layers of protein structures and their replacements with Gly should not contribute to reduced conformational stability, apart from structural damages arising from the lack of critical ion pair formation. On the other side, they have often a critical role in protein-protein interactions and assembly. Thus, the Gly-substitution of Arg, Asp and Glu, weakening the protein-protein recognition process, can be responsible for pathogenicity. Among amino acids with charged side chains, Arg/Gly replacements seem to be particularly pathogenic, possibly for the observed Arg critical involvement in protein-nucleic acid interactions [17].

**Figure 1:**
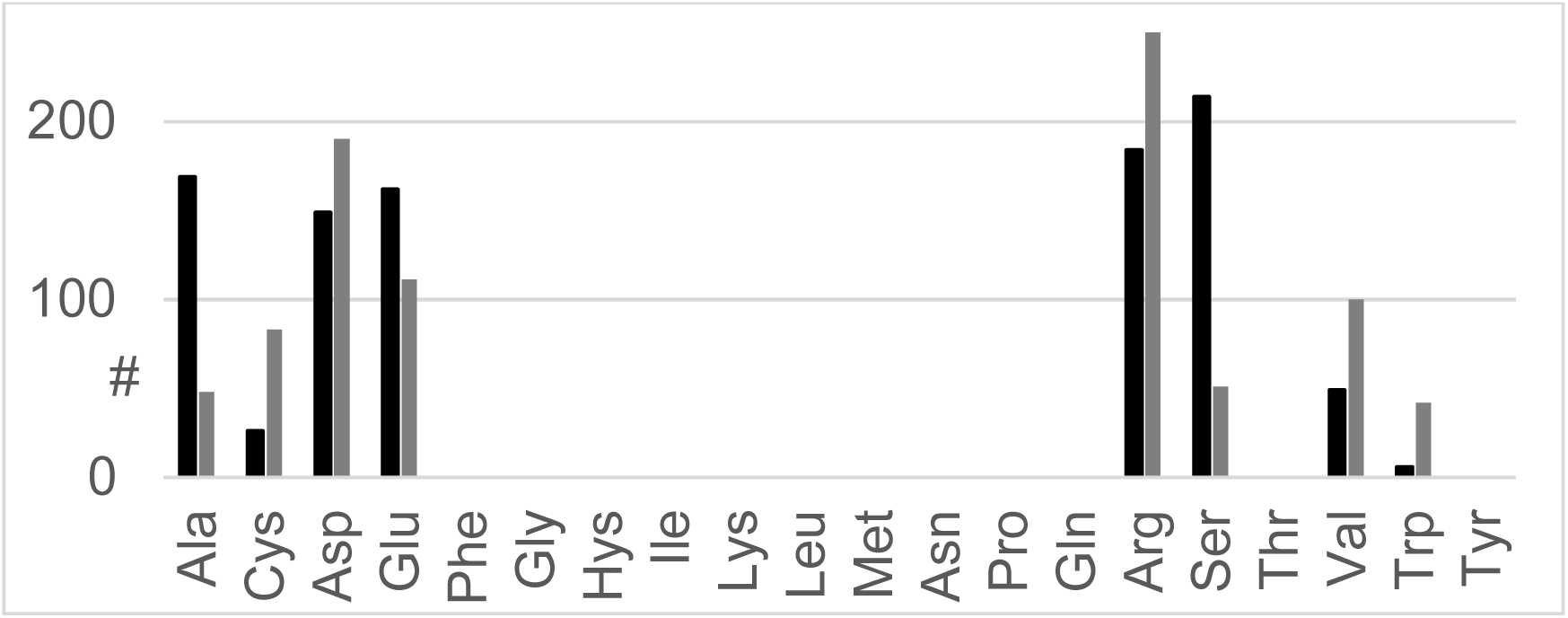
Distribution of X/Gly replacements referenced by ClinVar. Black and gray histograms respectively refer to Gly-substitution in benign and pathogenic missense mutations.

Then, we focused our attention on the 875 X/Gly pathogenic mutations to find all the features that can be responsible for the corresponding Mendelian disorders at an atomic level. It is a prerequisite for such investigation the possibility to assign a reliable X/Gly mutation topology for all the proteins of our data set. Accordingly, by taking into account only proteins that are deposited in PDB, that is with experimentally resolved structures, we could deal with 357 X/Gly mutated proteins for further analysis. However, the Xn that ClinVar delineates as a pathogenic mutation must be included in the protein PDB file, with the nth protein sequence position occupied by the amino acid to be replaced by Gly. We had to consider also the few cases where X/Gly mutations were so relevant that mutant structures have been characterized before or together with the wild-type protein. Taking into account these two limiting conditions, we had to remove 102 ClinVar items, getting a final data set containing 255 elements that we have structurally analyzed.

Then, we used *Gly-pipe*, a software that we have recently developed to check whether Gly mutations occurring close to the protein surface can cause changes in its shape [18]. This software compares native and Gly-mutated structures to find new pockets close to the mutation site. Thus, *Gly-pipe*, once applied to our data set of 255 X/Gly elements, indicates that only 96 mutations are located on protein surfaces, making them potentially responsible for functional modulations due to solvent dynamics changes. *Gly-pipe* is a machine learning procedure that includes the Pockdrug algorithm [19] to evaluate the druggability score of new or enlarged pockets determined by X/Gly replacements. We restricted our analysis to those protein surface changes that can induce possible druggable sites, *i*.*e*. with DS > 0.5 according to a previous analysis [20]. Thus, only 19 proteins of our data set have surface pockets induced by X/Gly substitutions with druggability scores suitable for ligand binding. Table 1 shows the 19 Mendelian disorders that are associated with Gly-mutations potentially altering protein surface dynamics and, hence, the biological function.

**Table 1.**

| **Mendelian Disorder** | **OMIM Code** | **Protein PDB ID** | **X<sub>n</sub>Gly** | **DS<sup>a</sup>** |
| --- | --- | --- | --- | --- |
| Permanent neonatal diabetes mellitus 3 | #618857 | 6C3O | Val86Gly | 0.670 |
| Ataxia-oculomotor apraxia type 1 | #208920 | 6CVQ | Val263Gly | 0.745 |
| Early infantile epileptic encephalopathy 2 | #300672 | 4BGQ | Trp195Gly | 0.703 |
| Deafness, autosomal dominant 9 | #601369 | 1JBI | Val66Gly | 0.674 |
| Mirror movements 1 | #157600 | 4PLO | Val793Gly | 0.544 |
| Congenital myasthenic syndrome 13 | #614750 | 6FWZ | Val264Gly | 0.729 |
| Waardenburg syndrome, type 4A | #277580 | 6IGK | Ala183Gly | 0.518 |
| Craniofrontonasal syndrome | #304110 | 6P7S | Trp37Gly | 0.513 |
| Alkaptonuria | #203500 | 1EYB | Val300Gly | 0.519 |
| Charcot-Marie-Tooth disease, axonal, type 2S | #616155 | 4B3F | Val373Gly | 0.563 |
| Long QT syndrome 2 | #613688 | 4HQA | Glu58Gly | 0.719 |
| Gastrointestinal stromal tumor | #606764 | 2VIF | Val569Gly | 0.519 |
| Multiple endocrine neoplasia, type 1 | #131100 | 6O5I | Glu45Gly | 0.666 |
| Lynch syndrome I | #120435 | 3THX | Val163Gly | 0.622 |
| Neurodevelopmental disorder with behavioral abnormalities, absent speech, and hypotonia | #618718 | 3ZYG | Trp107Gly | 0.600 |
| Amyotrophic lateral sclerosis 18 | #614808 | 4X1L | Cys71Gly | 0.750 |
| Noonan syndrome 4 | #610733 | 3KSY | Arg552Gly | 0.560 |
| Spastic paraplegia 4, autosomal dominant | #182601 | 5Z6R | Asp444Gly | 0.724 |
| Pheochromocytoma | #171300 | 4AJY | Trp117Gly<br>Arg107Gly<br>Ser111Gly | 0.758<br>0.674<br>0.674 |

a) Druggability scores were determined with *Gly-pipe* (9).

## 4. Discussion

From the initial 910 proteins that ClinVar reports to have X/Gly pathogenic mutations, we have ended up with only 19 cases fulfilling all our limiting conditions. Thus, in these 19 cases Gly-mutants present surface pockets that could be the target of small molecules to restore their original surface shape, and, therefore, protein dynamics and function.

Among the latter 19 proteins related to Mendelian disorders, we report here the discussion of six cases that reflect how differently Gly-mutants can determine pathogenicity.

The first example is given by the Von Hippel-Lindau protein (VHLP). Fig. 2 shows the structural effect of Arg107Gly mutation: here the Gly substitution causes a remarkable enlargement of the VHLP binding site towards hypoxia-inducible factor 1-alpha [21], causing a drastic change in the properties of their interaction. This feature may give a relevant contribution to understanding the carcinogenesis of pheochromocytoma (OMIM code 171300), a rare neuroendocrine tumor of the adrenal medulla. The design of ligands that can repair the structural damage due to the Arg107Gly mutation may represent a possible strategy to cure this Mendelian disorder.

**Figure 2:**
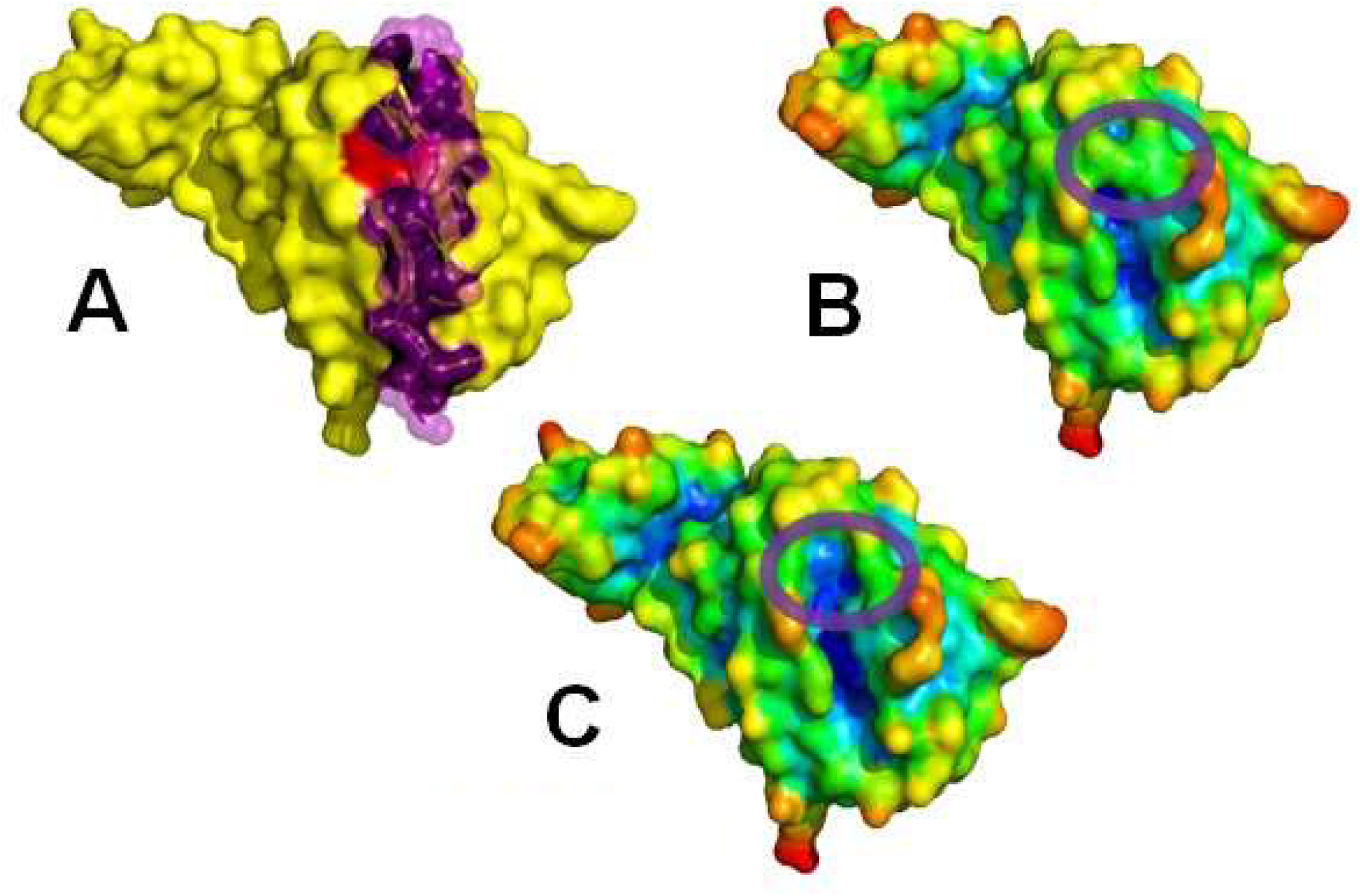
The HIF-1α interaction domain of VHL protein. Using the PDB 4AJY structure, in A is represented the VHL binding to a HIF-1α fragment, shown in purple transparent spheres. In B and C the wild type and the Arg107Gly-mutated VHL domains are respectively shown; PyMOL generated surfaces are colored according to protein atom depths; purple circles highlight the mutation site.

In the second example, we considered the SH2 domain of the suppressor of cytokine signaling 6 which includes the c-KIT binding site [22]. We can observe how the Val569Gly mutation in the SH2 domain modifies the structural features of c-KIT binding site, see Fig. 3. In this case, Gly-mutation has been associated with the insurgence of gastrointestinal stromal tumors (OMIM code 606764). As in the case of VHLP, again the Gly-mutation causes a modification of the protein-protein interaction pattern that a suitable ligand could restore.

**Figure 3:**
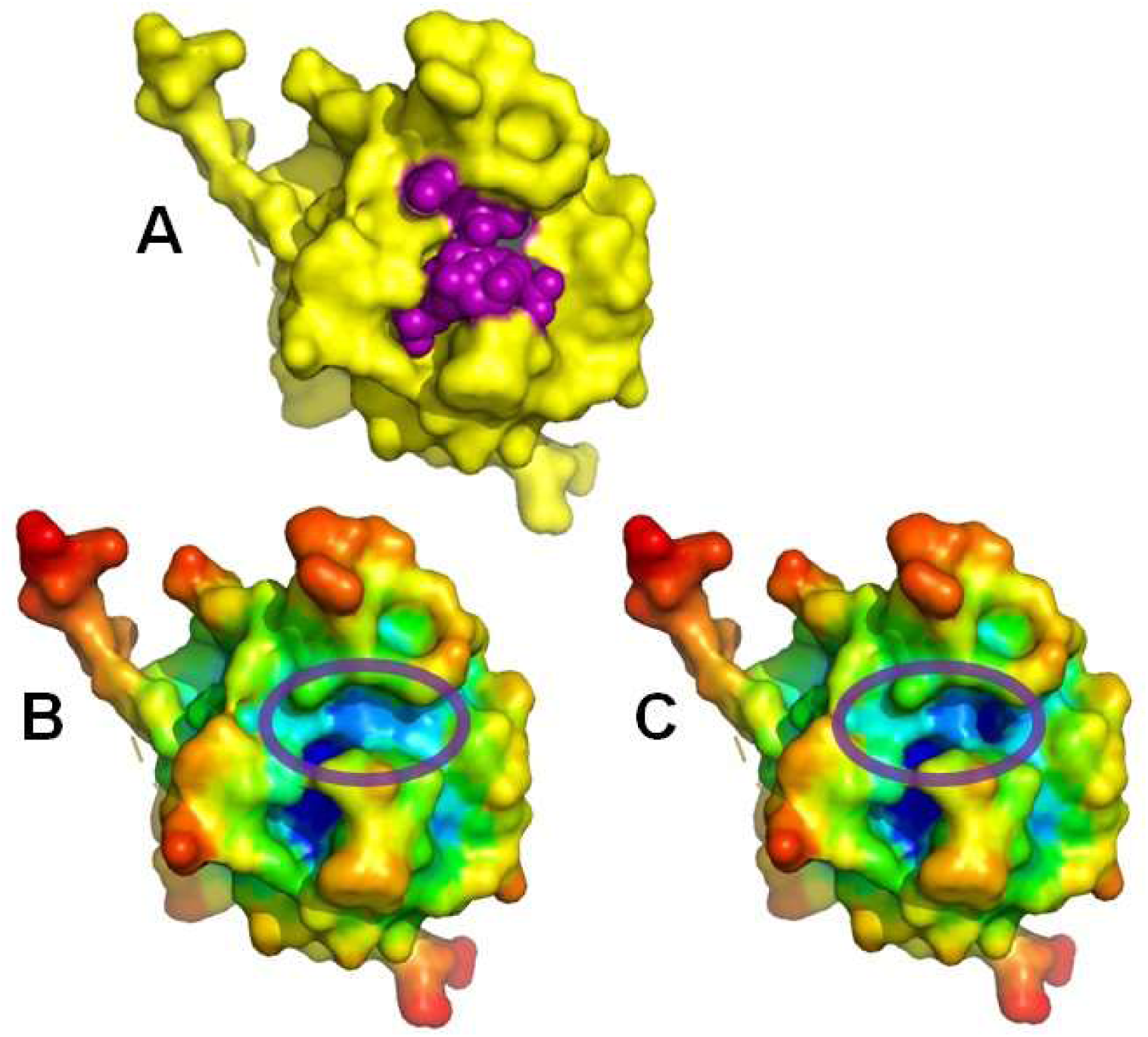
The SH2 domain of suppressor of cytokine signalling 6 bound to a c-KIT substrate fragment (PDB code 2VIF). In A, the c-KIT substrate, a 11 amino acid peptide, is shown in purple spheres. In B and C the wild type and the Val569Gly-mutated SH2 domains are respectively shown; PyMOL generated surfaces are colored according to protein atom depths; purple circles highlight the mutation site.

Among the 19 pathologies associated with Gly-mutations, as a third example, we have selected the autosomal dominant spastic paraplegia 4 (OMIM code 1826601). In this case, we analyzed the PDB structure of the AAA domains of spastin where the pathogenic Asp444Gly mutation occurs. As shown in Fig. 4, the Gly-mutation is located very near to the protein substrate ATP. Structural inspection of the surface cavity induced by the latter mutation shows the formation of a highly hydrophobic pocket, suitable for small molecule binding. It follows that AAA subunit interaction and/or its ATP binding, both fundamental processes for physiological activity [23] could be altered. Long QT Syndrome Type 2 (LQT2) with the OMIM code 613688, is the fourth Gly-mutation associated pathology that we have considered. This pathology has been ascribed to a reduced potassium current flowing through the hERG channel encoded by the KCNH2 gene [24]. The crystal structure of the PAS domain of the KCNH2 potassium channel [25], comprising the pathological Glu58Gly mutation site is available. Hence, we have compared the structures of this PAS domain in the absence and in the presence of the Gly-mutation, as shown in Fig. 5. The lack of a charged amino acid in the PAS domain surface exposes also a new hydrophobic pocket formed by Phe68 and Val59, a feature that may be reasonably associated with LQT2.

**Figure 4:**
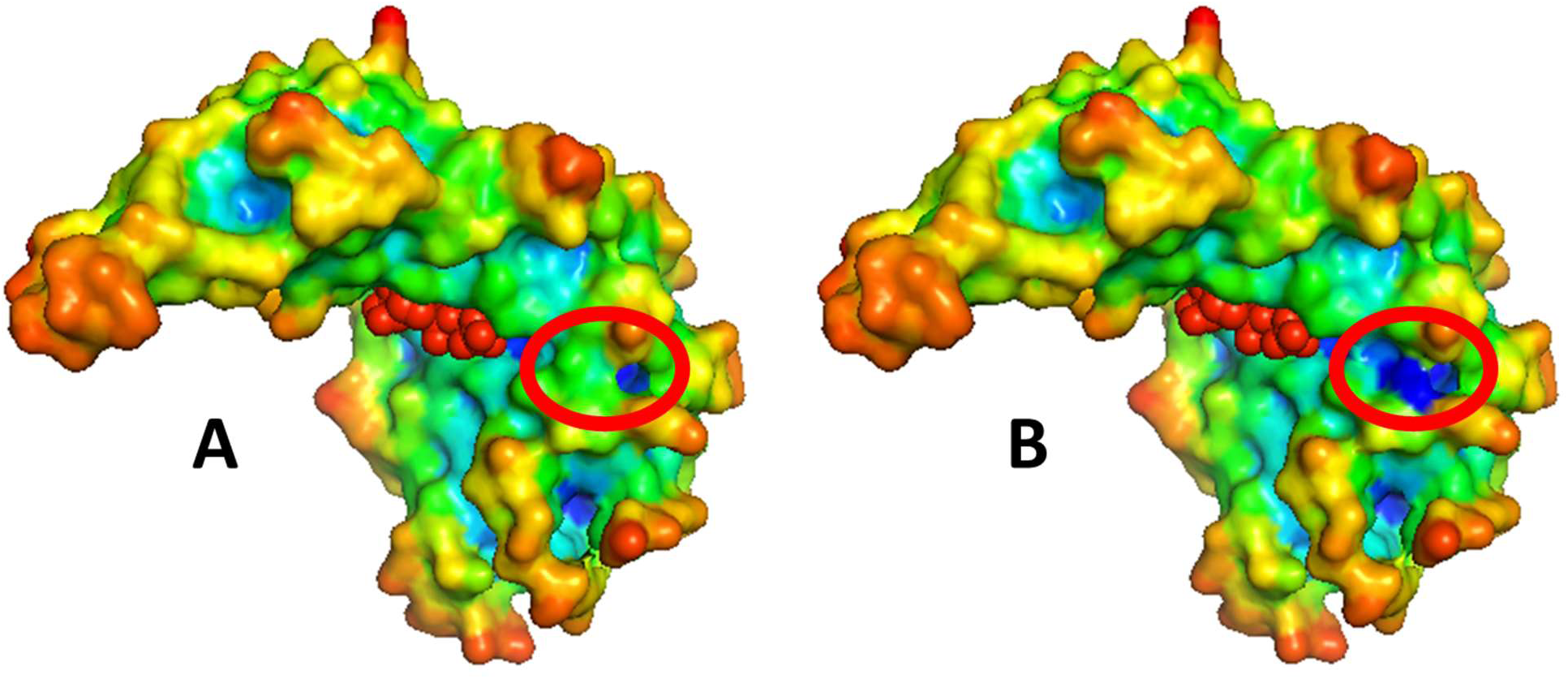
The AAA domain of spastin (PDB code 5Z6R). PyMOL surface representation of the protein domain complexed with ATP, shown in red spheres; A and B show respectively wild type and Asp444Gly mutant. Protein surfaces are colored according to protein atom depths; red circles highlight the mutation site.

**Figure 5:**
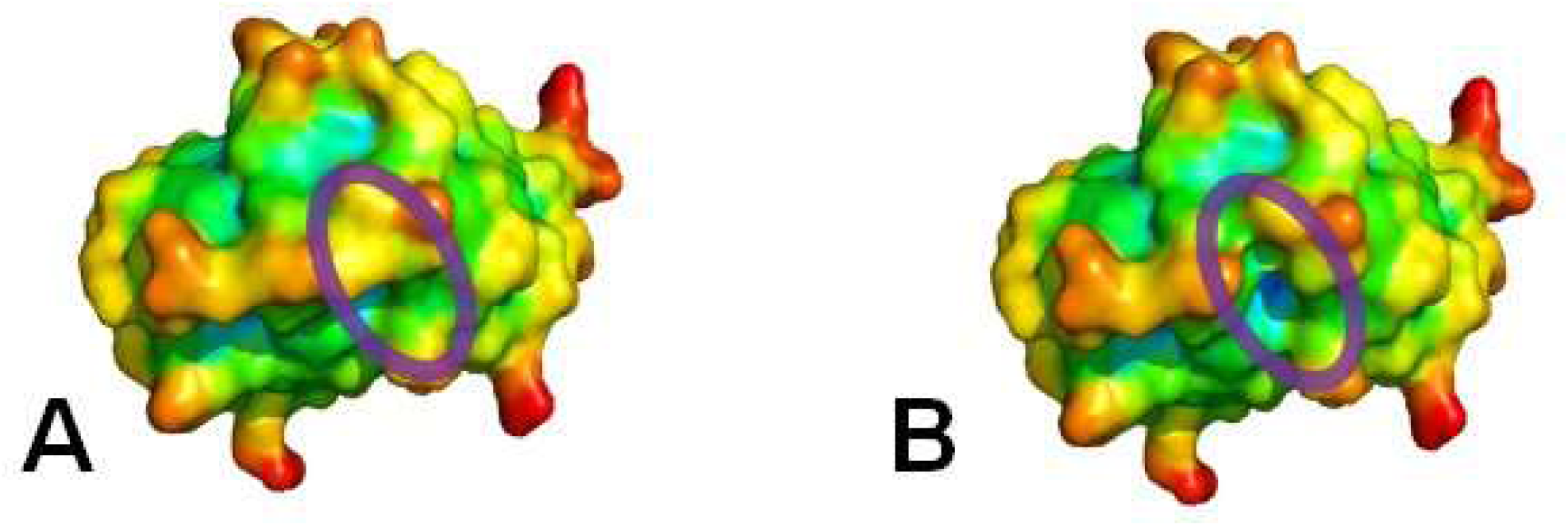
The PAS domain of KCNH2 potassium channel (PDB code 4HQA). A and B show respectively wild type and Val569Gly mutant; protein surfaces were generated by PyMOL and colored according to protein atom depths; purple circles highlight the mutation site.

Data reported in Fig. 1, clearly indicate that Trp/Gly replacements are the most pathogenic ones among all the eight possibilities given by a single nucleotide change in the genetic codon. This finding is largely expectable, as amino acids with the biggest side chain are replaced by the smallest ones. Furthermore, substitutions of Trp, due to its variable topology [26] can alter the protein surface shape in the case that it is located in a peripheral moiety, see the two examples shown in Fig. 6. The Trp117Gly mutant of the VHLP, together with the above-discussed Arg107/Gly and the Ser111Gly mutants, contributes to the occurrence of pheochromocytoma by modifying the VHLP surface dynamics. Mutations in the human CDKL5 kinase domain have been considered contributing causes of early infantile epileptic encephalopathy 2 [27]. The Trp195Gly, modifying the protein surface shape, see Fig. 6C and 6D, can represent the molecular origin of this Mendelian disorder. However, Trp more frequently occupies inner positions that are fundamental for the folding nucleus formation and, hence, a Trp/Gly substitution can severely compromise the protein structural stability.

**Figure 6:**
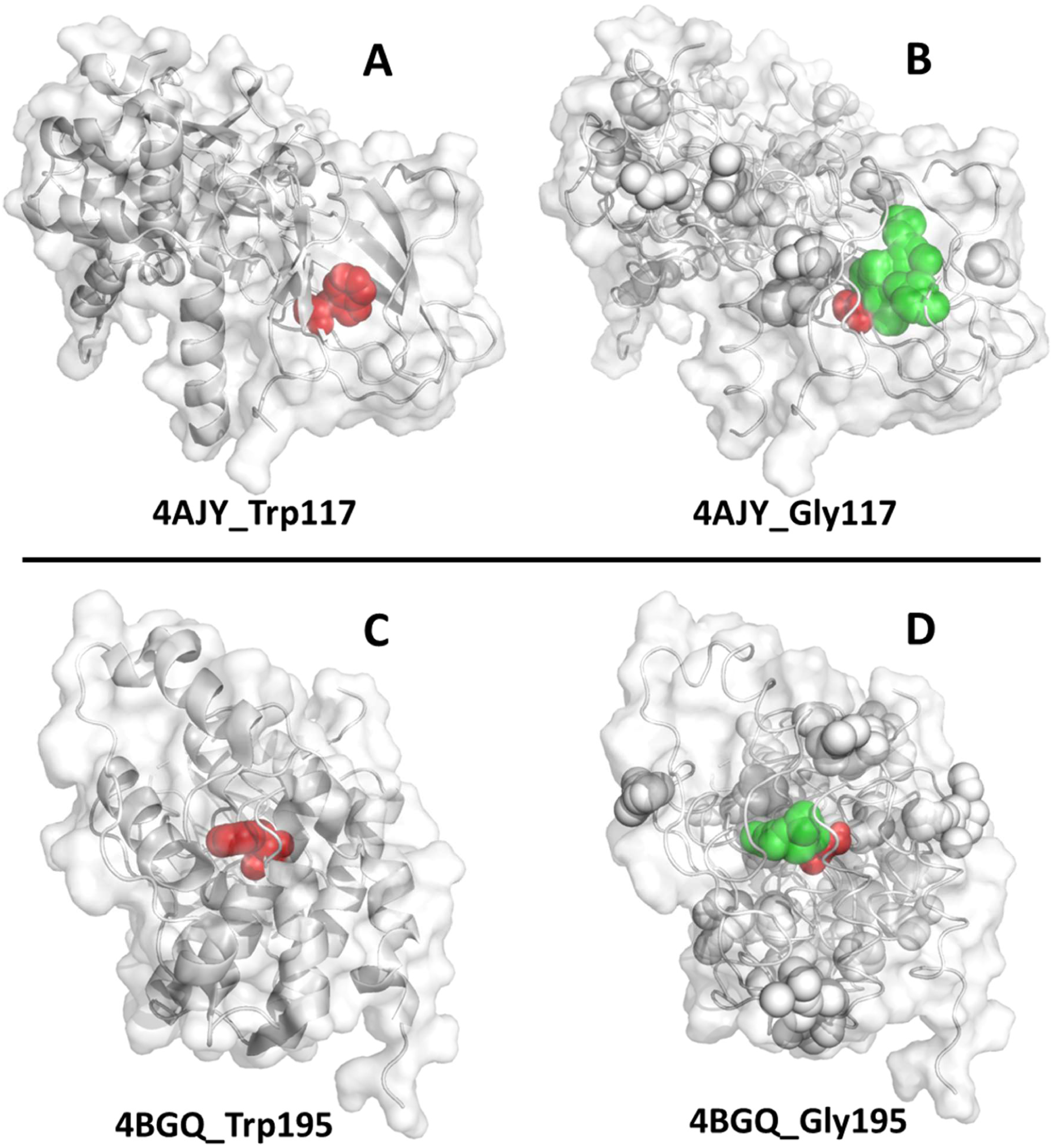
The structural consequence of Trp/Gly replacements. In A and B, the wild type and the Trp117Gly mutant of the von Hippel-Lindau protein are respectively shown. In C and D, the same comparison between wild type and Trp195Gly mutant is shown for the human CDKL5 kinase domain. Trp and Gly atoms are colored in red spheres; alpha spheres, generated by Fpocket, are shown in B and D; green alpha spheres highlight the pocket induced by Trp/Gly substitutions

We did not perform any virtual screening of ligands that could be suitable to bind pockets arising from Gly mutations not to add predictions to our predictive approach. The virtual screening procedure, indeed, would require preliminary qualitative and quantitative characterizations of geometry and lifetime of the open form for the predicted Gly-induced pocket, currently under investigation in our laboratory through Molecular Dynamics simulations.

## 5. Conclusions

We have proven, even though in a limited number of cases, that Gly-mutations can be responsible for the formation of new pockets on the protein surface if some conditions are fulfilled, as in the case that amino acids involved in the replacement bear bulky side chains and are located near to the protein surface [9]. This feature can be exploited to design Gly-mutants for proteins whose activity we like to control through suitable ligands. In the present report, we explored how evolution designed Gly-mutants as witnessed in the human genome by ClinVar. ClinVar, as of July 4, 2022, classified 959 benign and 875 pathological missense Gly-mutations, offering a large repertoire for understanding the way these mutations can cause Mendelian disorders. By crossing ClinVar and PDB data banks, the structural feature that originates a Mendelian disorder can be analyzed and, in case the Gly-mutation determines the formation of a potential new binding site, remediation could be found by discovering ligands that shield the latter binding site, fixing a Mendelian disorder. It must be underlined that a severe bottleneck for the present investigation was due to the lack of structural information, reducing the 875 proteins with pathological Gly-mutations to 225 structurally characterized systems. The continuous progress in protein structure predictions, like the ones recently proposed [28-30], will solve in part the problem. In general, AI will yield more and more powerful tools to predict which and how Mendelian disorders can be cured. Furthermore, the AI era opens new perspectives to drug design [31] and protein-ligand simulations [32]. The possibility of very fast tailoring of ligands for Gly-induced pockets through molecular graph generation with graph neural networks is just a very recent example of AI-related advancement [33], whose products can also be automatically evaluated with an AI predictor of side-effects [34]. Thus, our structural Bioinformatics survey on the human pathogenic variant database suggests a fast rational selection of those Mendelian disorders where X/Gly-mutations determine damages that can be repaired simply by swallowing a pill: a very powerful shortcut to restrict all the experimental procedures that are needed to achieve disease remediation.

As a final remark, we like to underline that the present investigation shed light, at a molecular level, on the molecular mechanisms of Mendelian disorders due to Gly-mutations, offering a powerful aid to select new therapeutic strategies.

## Abbreviations

SADIC: Simple Atom Depth Index Calculator
Gly: glycine
Arg: arginine
VHLP: Von Hippel-Lindau protein

## Acknowledgments

Thanks are due to Prof. P. A. Temussi for helpful discussions and to Valentina Pacciani for technical assistance. This research did not receive any specific grant from funding agencies in the public, commercial, or not-for-profit sectors.

## Notes

### Competing Interest Statement

The authors have declared no competing interest.

